# Therapeutic restoration of synaptic architecture, retinal and visual function, and prevention of retinal degeneration in a mouse model of retinal dystrophy

**DOI:** 10.64898/2026.06.23.733438

**Authors:** Nazarul Hasan, Mattia D. Paolo, Maureen A. McCall, Ronald G. Gregg

## Abstract

Vision depends on the transfer of photoreceptor signals through the retina and then to many CNS visual nuclei. While the most common inherited retinal diseases (IRDs) involve defects in rod and/or cone function, another group (referred to as congenital stationary night blindness (CSNB)) results from defects in glutamate release from photoreceptors, or conversion of the glutamatergic signal in bipolar cells. One example results from mutations in the *CACNA2D4* gene, which encodes a subunit of the voltage-gated calcium channel that is critical for glutamate release from both rod and cone photoreceptors. Mutations in *CACNA2D4* result in a range of phenotypes in human patients, from incomplete CSNB to rod-cone dystrophy. In the *CACNA2D4* knockout mouse (*α2δ4^-/-^*), there is slow photoreceptor degeneration, the photoreceptor-to-bipolar cell synapse is disorganized, and the retina lacks scotopic and photopic full-field electroretinogram b-waves; this also results in low visual acuity. Using adult *α2δ4^-/-^*mice, we show that recombinant adeno-associated virus (rAAV)-mediated gene therapy directed to rod photoreceptors prevents rod degeneration, restores synaptic organization, retinal function, and improves visual acuity under both light- and dark-adapted conditions. This rescue was maintained for up to 14 months post-treatment. Together, our results demonstrate that synaptic structure and function can be restored in the mature mouse retina in a model of complete synaptic disorganization. The results highlight the neuroprotective potential of targeting synaptic organizing proteins in retinal gene therapy.

## INTRODUCTION

Vision begins in the retina when rod and cone photoreceptors detect light and convert it into an electrical signal that alters glutamate release from their terminals. The photoreceptors’ terminals form synapses in the outer plexiform layer (OPL) with bipolar cells (BCs), establishing a vertical signaling pathway that is modulated by lateral feedback from horizontal cells (HCs). Photoreceptor signaling to BCs and HCs is glutamatergic, and release is mediated by the voltage-dependent calcium channel (Ca_v_1.4 VDCC), CACNA1F ^1^. Rod photoreceptors support vision in low-light conditions, whereas cone photoreceptors support vision in bright light. The primary output of rods is to ON rod BCs, whereas cones connect to two general types of ON and OFF cone BCs. All ON-type BCs detect changes in glutamate release via the metabotropic glutamate receptor 6 (mGluR6), which modulates the opening of the TRPM1 cation channel. OFF BCs detect changes in glutamate release via AMPA/kainate ionotropic glutamate receptors (see review ^2^).

The photoreceptor-to-ON bipolar synapse is specialized and associated with the presence of presynaptic ribbons. In the mouse, this synapse develops and matures prior to eye opening (∼postnatal day 10). Specifically, it is organized as an invagination of the ON rod or cone BC dendrites and HC axons into the photoreceptor terminal (rod or cone). Because the structural integrity of this synapse is required for normal visual function, mutations in genes whose proteins make up rod to rod BC synaptic complexes cause vision defects, including complete and incomplete congenital stationary night blindness (cCSNB, iCSNB), retinal dystrophies, and retinitis pigmentosa ^1,3^.

In this report, we focus on one subunit of the photoreceptor Ca_v_1.4 VDCC, CACNA2D4 (α2δ4), which is critical for glutamate release and required for calcium channel trafficking, localization, and gating properties ^4^. Patients with mutations in *CACNA2D4*, have a spectrum of phenotypes, including autosomal recessive (AR) cone dystrophy, retinitis pigmentosa, cone—rod dystrophy, and stable mild dysfunction ^3,5–7^. Two mouse models in which *Cacna2d4* was knocked out, have an iCSNB phenotype that includes: absent or greatly reduced electroretinogram b-waves, disrupted photoreceptor to bipolar cell synaptic architecture, and reduced visual sensitivity ^4,8,9^, and in aged mice, a rod-cone dystrophy develops with retinal degeneration^4^. These findings establish α2δ4 as an essential organizer of ribbon synapse assembly and function.

Recombinant adeno-associated virus (rAAV) vectors have emerged as a powerful tool for delivering therapeutics to photoreceptors to restore vision in patients with RPE65-associated Leber congenital amaurosis ^10,11^. However, it remains unknown whether rAAV-mediated gene delivery to photoreceptors can restore protein expression, synaptic architecture, and visual function when synapse organization is completely disrupted. We addressed whether we could restore structure and function in the mature retina. We delivered rAAV encoding wildtype (WT) α2δ4 to rod photoreceptors of *α2δ4^-/-^* adult mice. Our results show that restoration of α2δ4 expression restores rod function, normal synaptic organization, and visual acuity in adult mice.

## RESULTS

The absence of the α2δ4 subunit of the VDCCs prohibits the proper assembly of the Ca_v_1.4 L-type VDCCs at photoreceptor synapses. The retinas of *α2δ4^-/-^* mice exhibit extensive disorganization of the synaptic architecture between photoreceptors and bipolar/horizontal cells, resulting in the collapse of the outer plexiform layer (OPL), synapse disruption, photoreceptor axon retraction, an abnormal ffERG, reduced light sensitivity, and ultimately retinal degeneration ^4,8,9^. We evaluated whether restoring expression of the missing α2δ4 protein via rAAV gene therapy could restore the Ca_v_1.4 VDCC as well as synapse structure and retinal/visual function in the mature retina, where plasticity is less likely to play a role. We reintroduced α2δ4 into the retinas of P35 *α2δ4^-/-,^* hereafter called KO, mice via subretinal injection of rAAV RHO::α2δ4. Because the loss of α2δ4 affects the molecular organization of rod synapses more severely than cone synapses ^4^, we focused on delivering α2δ4 to rods (and the rod-to-rod bipolar synapse) using a well characterized rod specific promoter ^12,13^. A schematic of the rAAV construct, rAAV RHO::α2δ4, and the experimental design are shown in Fig. 1B We injected 1 x 10^10^ rAAV RHO::α2δ4 viral genomes subretinally into mature ∼P35 day old KO retinas and assessed retinal and cortical function, as well as retinal cellular morphology (Fig. 1B). Injection of GRK1::GFP into KO retinas served as a control. Hereafter the groups are referred to as WT (C57BL/6), Control (WT treated with rAAV GRK1::GFP), KO (*α2δ4^-/-^*), KO Control (*α2δ4^-/-^* treated with rAAV GRK1::GFP), KO-Treated (*α2δ4^-/-^* treated with rAAV RHO::α2δ4).

**Figure 1.**
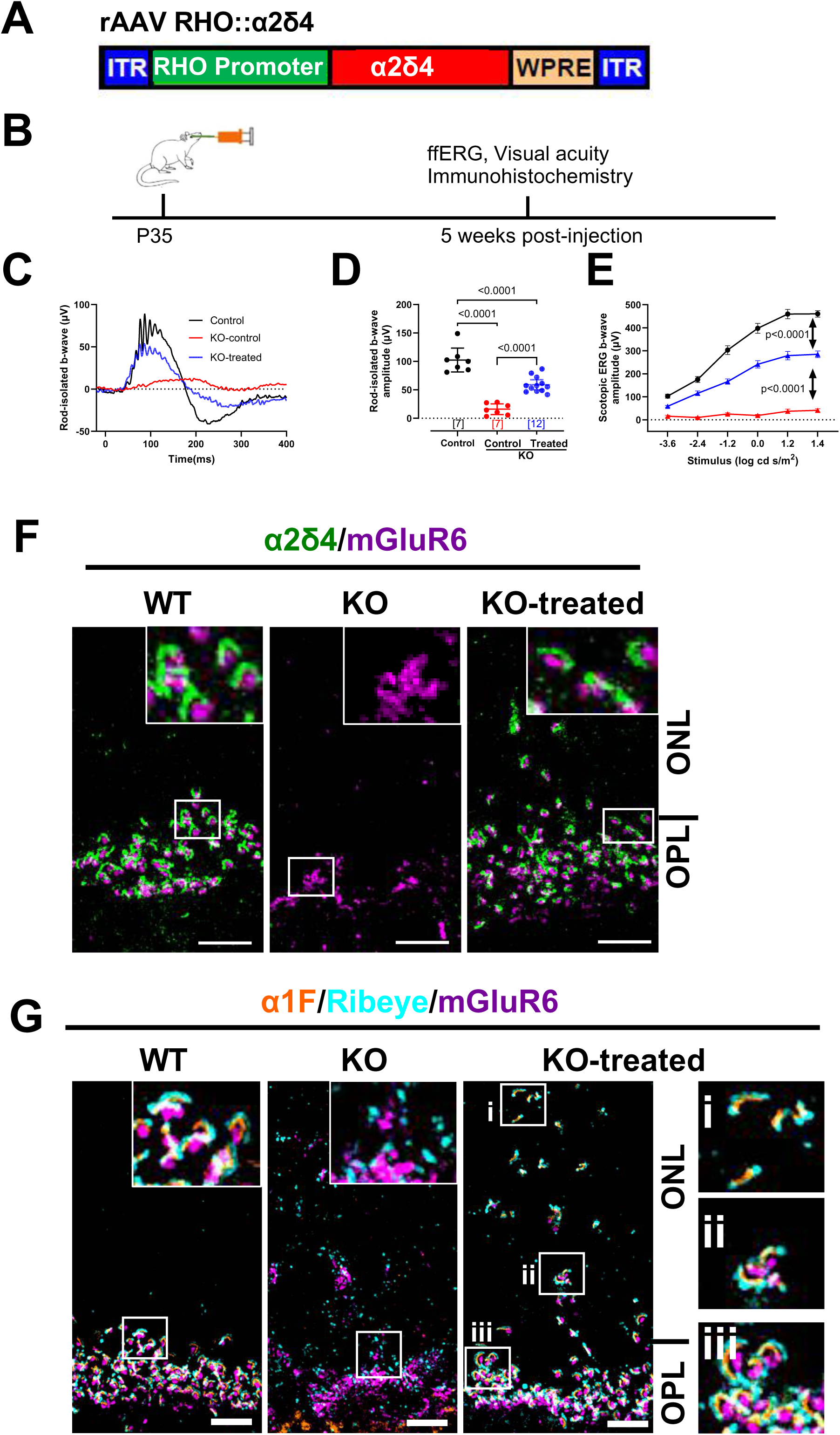
Expression of α2δ4 in rods of *α2δ4^-/-^*retinas restores function. (A). Schematic diagram of the rAAV RHO::α2δ4 vector, showing the RHO promoter and coding region for the α2δ4 protein. (B). Time course of treatment and analysis. (C). Scotopic rod-isolated (flash intensity = -3.6 log cd.s/m^2^) ffERG responses for a single mouse: Control (black, WT treated with rAAV GRK1::GFP), KO-control (red: *α2δ4^-/-^* treated with rAAV GRK1::GFP), and KO-treated (blue; *α2δ4^-/-^* treated with rAAV RHO::α2δ4). (D). Scotopic rod-isolated ffERG amplitudes from multiple mice (numbers indicated below X axis). (E). Mean scotopic ERG b-wave amplitudes (± SEM) at multiple flash intensities for mice shown in D. (F). Representative merged images of transverse retinal sections of WT, KO and KO-treated retinas stained with antibodies to α2δ4 (green) and mGluR6 (magenta). Insets show higher-power images to demonstrate the characteristic horseshoe staining pattern of α2δ4 surrounding mGluR6 puncta in WT and KO-treated retina that is absent in the KO. (G). Merged images of transverse WT, KO and KO-treated retina sections stained with antibodies to α1F (orange), ribeye (cyan), and mGluR6 (magenta). Insets WT and KO show higher-power images of boxed regions to demonstrate the close association of α1F and ribeye. (Gi-iii) High power images of regions boxed in KO-treated retina illustrating the different composition of some restored horseshoe shaped elements. Images are representative of those from at least 5 retinas. Scale bars = 5µm. Statistics: D, ANOVA, and E two-way ANOVA with adjustment for multiple comparisons. p≤ 0.05 is considered significant.

### Expression of α2δ4 in rods of KO restores signal transmission to bipolar cells

The abnormal synaptic transmission between photoreceptors and bipolar cells in KO retina can be detected using the full-field electroretinogram (ERG; ^14^). At P35 (the time of treatment), there is a small but measurable scotopic b-wave in KO retinas. We compared the rod-isolated scotopic ffERG responses in KO-treated, KO controls, and Control mice and measured the rod-isolated b-wave 5 weeks post-injection (wpi). Waveforms for single animals (Fig. 1C) and amplitudes for multiple mice (Fig. 1D) show that the rod isolated b-wave was restored in KO-treated mice, but not in KO-controls. The amplitude of the rod-isolated ERG b-wave in the KO-treated mice was restored to ∼57.6% of Control (Fig. 1D,E; n = 12; p = <0.0001) and was significantly increased compared to KO-controls. This result was also reflected in significantly larger b-wave responses across all flash intensities (Fig. 1E).

### Expression of α2δ4 in KO rods restores VDCC channel expression and rod synaptic structure

In the KO mouse retina, both the VDCC and the normal invaginating synapse in the rod spherule are absent ^4,8^. Instead, rod axons retract into the ONL, and rod BC dendrites and horizontal cell (HC) axons processes extend into the ONL, appearing to follow the rod axons, which collapses the OPL. This phenotype is consistent with other models in which Ca_v_1.4 VDCC expression and/or function is altered ^15^. We first examined the expression pattern of α2δ4 relative to the post synaptic marker, mGluR6, in WT, KO, and KO-treated retinas (Fig. 1F). Our images show the characteristic WT horseshoe shaped staining pattern of α2δ4 is lost in the KO and restored in the KO-treated retinas. Further, mGluR6 puncta present and inside the α2δ4 horseshoes of WT are lost in the KO and restored in the KO-treated retinas (Fig. 1F). To evaluate synapse structure further, we triple-stained to co-localize α1F, the main pore-forming subunit of the Ca_v_1.4 VDCCs, with ribeye, and mGluR6. In WT retina, α1F is located near the ribbon, marked by ribeye, and both form a characteristic horseshoe pattern, with mGluR6 puncta located inside the horseshoe (Fig. 1F,G). This pattern is completely disrupted in the KO retina (Fig. 1F, and ^15^). No horseshoe shaped α1F or ribeye positive profiles are found, and ribeye expression appears punctate, similar to WT mGluR6. There is punctate expression in both the OPL, and throughout the ONL (Fig. 1G). In contrast, treatment with rAAV RHO::α2δ4 in KO mice restores the WT staining pattern in the OPL, establishing that the Ca_v_1.4 VDCC, marked by α1F expression, is trafficked correctly. Surprisingly, in KO-treated retinas, horseshoe-shaped α1F and ribeye positive profiles are found in the ONL, likely identifying ectopic synapses (Fig 1G). The expression of α2δ4 also restores expression of other components of the rod synapse, including GPR179, TRPM1, RGS11, LRIT3, PSD95, and Pikachurin (Figures S1, S2).

To further define the level of rAAV RHO::α2δ4-mediated restoration of synaptic organization, we examined rod bipolar cell dendrites (PKCα, mGluR6) and horizontal cell processes (Calbindin) relative to ribeye and mGluR6 expression (Fig. 2, and ^4^). In the WT mouse retina, antibodies to PKCα stain the entire rod BC, and antibodies to calbindin stain horizontal cells as well as their processes, including the dendrites that form the invagination into the rod spherule (Fig. 2). Unfortunately, the calbindin and PKCα antibodies were both produced in rabbits and must be used separately.

**Figure 2.**
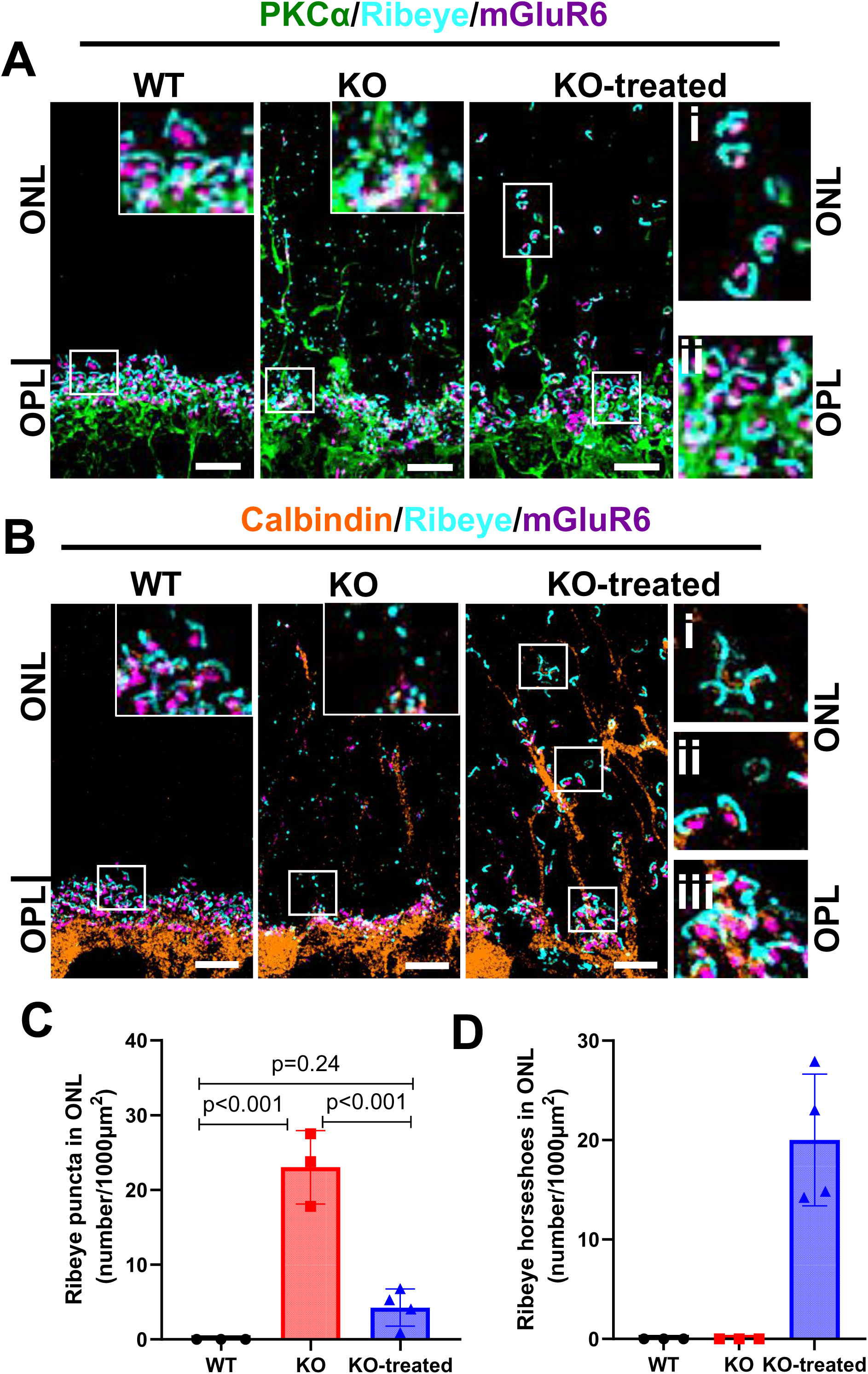
Expression of α2δ4 in *α2δ4^-/-^* retina restores rod synaptic organization. (**A**). Merged transverse retinal sections stained with antibodies to PKCα (green), ribeye (cyan), and mGluR6 (magenta) for WT, KO, and KO-treated retinas. Insets show high-power images of boxed areas and (Ai and ii) illustrated the pattern of these markers in the ONL and OPL of KO-treated retina. The WT PKCα bipolar cell profiles, normally restricted to the OPL, extend into the ONL in the KO and KO-treated retinas. (**B**). Merged images of retinal sections stained with antibodies to calbindin (orange), ribeye (cyan), and mGluR6 (magenta). Insets show high-power images of boxed areas in WT and KO, and Bi-iii illustrated the pattern of these markers in the OPL (Biii) and throughout the ONL(Bi and ii) in KO-treated retina. Note the absence of mGluR6 staining within the horseshoe in Bi, and the presence in Bii. Scale bar = 5µm. (**C**). Quantification of puncta and horseshoe shaped structures in the ONL of sections stained for ribeye and DAPI. One-way ANOVA with post-hoc correction for multiple testing. p≤ 0.05 is considered significant.

In KO retinas, PKCα positive rod BC dendrites and calbindin positive HC processes extend aberrantly into the ONL (Fig. 2, ^4,8^, where they associate with ribeye and mGluR6 puncta. In KO-treated retinas, these processes remain (Fig. 2 and S3). However, the synaptic arrangement of α2δ4 relative to ribeye and mGluR6 seen in WT does reform, regardless of location in the OPL or the ONL (Fig 2A). In the ONL, the number of ribeye puncta present in the KO and absent in WT are significantly reduced in the KO-treated ONL (Fig 2C). Also, the number of ribeye horseshoe-shaped structures absent in the KO ONL is significantly increased in the KO-treated ONL (Fig 2C).

The mGluR6 expression pattern in the treated KO retinas is variable, with two distinct patterns. In many instances, ribeye and mGluR6 show the expected juxtaposition in both the OPL and ONL (Fig. 2Aii and 2Aiii). In others, ribeye structures lack an opposing mGluR6 puncta (Fig. 2Bi), suggesting either the absence of the rod BC dendrite or mGluR6 expression below the detection threshold. In contrast, horseshoe-shaped structures in the ONL lacking mGluR6, showed calbindin labeling, confirming the presence of an HC process (Fig. 2Bi). Together, these data thus demonstrate that re-expression of α2δ4 in KO mice is sufficient to reform horseshoe shaped ribbon synapse structures, and that HC processes are required for this reformation, consistent with previous reports ^16^.

### Cone function is improved in the rAAV KO-treated retinas

We assumed that our well-characterized rod-specific promoter would selectively drive α2δ4 expression in rods, and were surprised to find a restored photopic ffERG b-wave in KO-treated mice, that is absent in KO-controls. In the KO-treated mice the photopic b-wave amplitude was restored to ∼50% of the Control (Fig 3A; individual mice; Fig 3B and C, all injected mice). Note that these data are from the same mice as shown in Fig. 1C. This result in the KO-treated retinas suggests the novel idea that the vector we used is also expressed in treated KO cones. To evaluate this, we double labeled retinal sections with PNA and antibodies to α2δ4 from WT, KO, and KO-treated retinas. In WT mice, α2δ4 expression is located immediately above the PNA label on cone terminals (ovals; Fig 3D and merge inset). In the KO mice, PNA staining remains in the absence of α2δ4 expression (Fig.3E) and cone structure appears somewhat disorganized, consistent with previous reports ^4,8^. In KO-treated retinas, α2δ4 re-expression is adjacent to PNA cone pedicles (Fig. 3F) and consistent with the photopic ffERG restoration.

**Figure 3.**
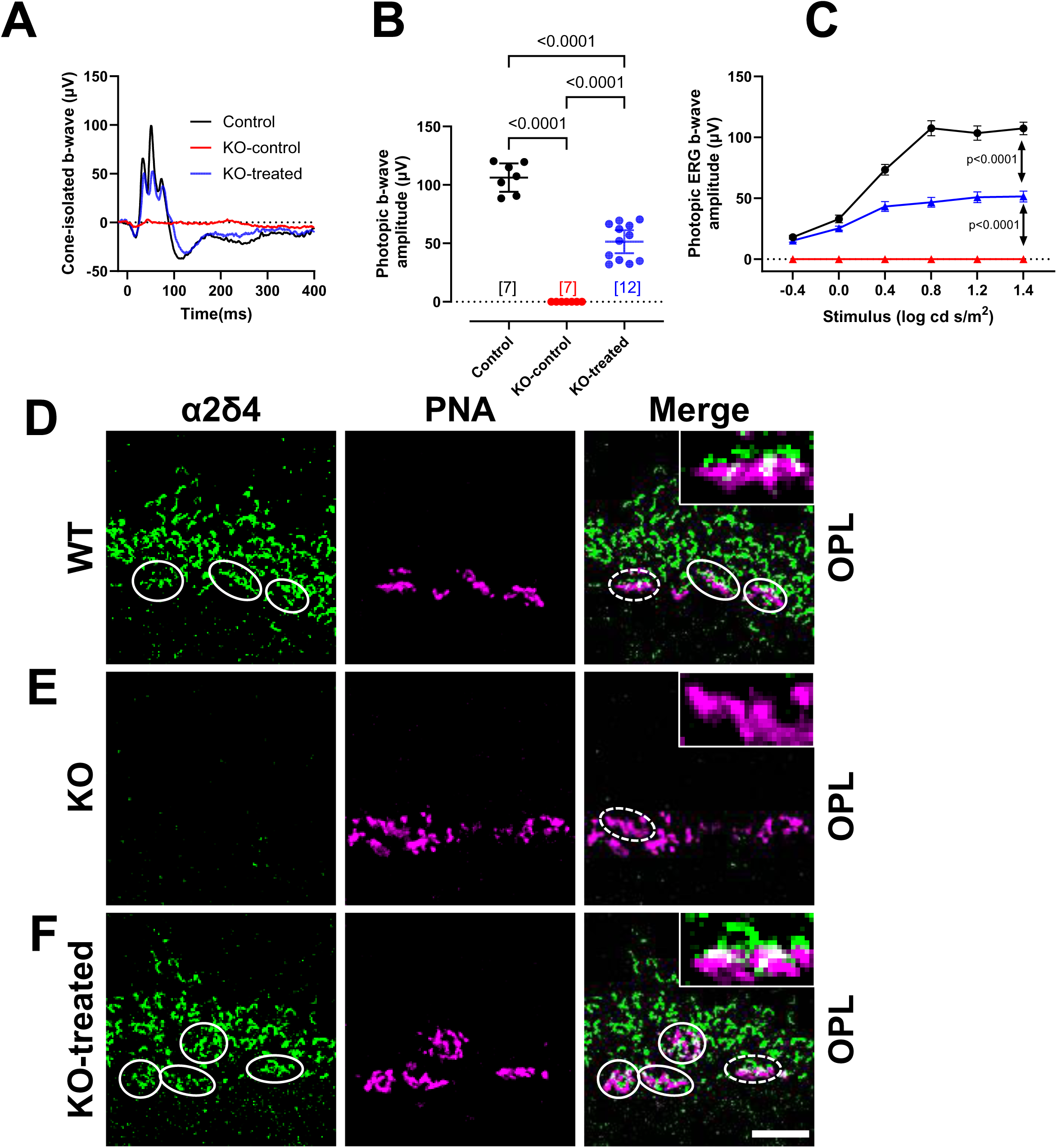
Expression of α2δ4 directed to KO rods restores cone function. (A). Photopic ffERG responses (flash intensity = 1.4 log cd.s/m^2^ on 20 cd/m² background) for single Control (black), KO-control (red), and KO-treated (blue) mice (same mice as in Fig. 1C). (B). Photopic ffERG amplitudes from all mice in each group (n indicated above the X axis). (C). Photopic ffERG b-wave amplitudes across multiple flash intensities for mice shown in B (and Fig. 1D). (**D-F**) Representative transverse retinal sections stained with PNA (magenta) and an antibody to α2δ4 (green), for WT (D), KO (E), and KO-treated (F). Insets show high-power images of boxed areas to illustrate staining of a single cone terminal. There is a close association of PNA and α2δ4 in WT that is absent in the KO and restored in treated KOs, consistent with α2δ4 expression in cones. Scale bars = 5µm. Statistics: B, one-way ANOVA, and C, two-way ANOVA with adjustment for multiple comparisons. p≤ 0.05 is considered significant.

### rAAV-mediated expression of α2δ4 in *α2δ4^-/-^* mice improves visual acuity

Our gene therapy approach should be durable, and to verify this assumption, we assessed visual acuity at 5 weeks post-injection (wpi), and ERGs and retinal structure at 3 and 14 months post-injection (mpi; Fig. 4A) to evaluate the effect of treatment on the reported slow degeneration in the KO retinas ^4,9^.

**Figure 4.**
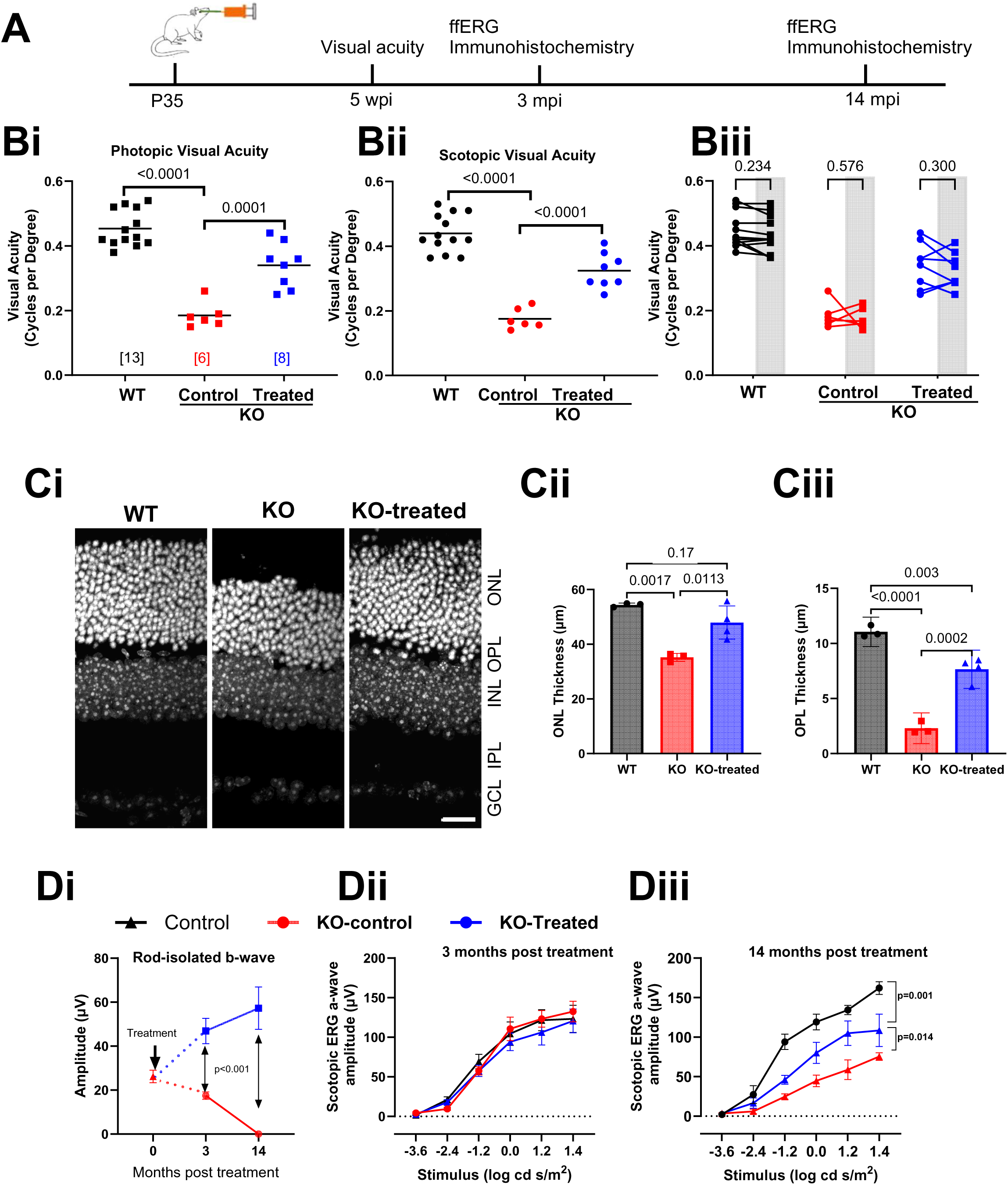
Expression of α2δ4 targeted to KO rods improves scotopic and photopic visual acuity and slows rod degeneration. (A). Schematic diagram of the rAAV RHO::α2δ4 vector, showing the RHO promoter and coding region for the α2δ4 protein. (B). Photopic (Bi) and scotopic (Bii) visual acuity of WT, KO-control, and KO-treated mice, respectively. (B**iii**) Visual acuity of each mouse under photopic and scotopic (gray bar) conditions. The same mice are plotted in all **A** graphs for clarity of comparisons. (Statistics: two-way repeated measures ANOVA analysis, with multiple comparisons correction. p≤ 0.05 is considered significant.) (C) Representative transverse retinal sections stained with DAPI (Ci) from age-matched WT, KO mice, and KO-treated at 14 months (14 mpi). (Bii) ONL thickness from age-matched WT, KO, and KO-treated mice at 14mpi. (Ciii) OPL thickness of mice in Bii. Statistics: one-way ANOVA with adjustment for multiple comparisons; p≤ 0.05 is considered significant. Scale bars = 5µm (D). Rod-isolated ffERG b-wave amplitudes (Di) for KO-control and KO-treated mice at baseline (time of treatment (P35)), 3 and 14 mpi. Data at P35 is from different mice than 3 and 14 mpi, (indicated by the dashed lines.) (Dii) Scotopic ffERG a-wave from 3mpi Control, KO-control and KO-treated at multiple flash intensities. (Diii) Scotopic ffERG a-wave at 14mpi from Control, KO-control and KO-treated at multiple flash intensities. (Statistics: Two-way repeated-measures analyses of the 3 and 14 mpi groups, adjusted for multiple comparisons. p≤ 0.05 is considered significant.)

To measure visual acuity, we used a two-alternative forced-choice visual discrimination task under light- and dark-adapted conditions at 5 wpi ^17,18^). The visual acuity of WT, KO-control, and KO-treated mice was first assessed under photopic conditions and subsequently after dark-adaptation. As we previously observed, visual acuity in WT mice is similar under both scotopic and photopic conditions. In KO-control mice, scotopic and photopic visual acuity were significantly reduced compared to WT (Fig 4Bi, Aii; WT 0.36 to 0.54 cycles per degree (cpd); p_adj_ = 0.234); KO, 0.14 to 0.26 cpd; respectively), consistent with their profound loss of retinal function (Fig 1C-E, ^4,8^). In KO-treated mice, scotopic and photopic visual acuity were significantly improved compared to untreated KO -control mice (0.25 to 0.41 cpd; p_adj_ = <0.0001 and 0.26 to 0.44; p_adj_ = 0.0001, respectively), and some KO-treated mice reached visual acuities comparable to WT. These results demonstrate that rAAV-mediated α2δ4 expression in photoreceptors of KO mice is sufficient to rescue both rod- and cone acuity to near WT levels.

### Durable gene therapy in KO-treated mice prevents retinal degeneration

To assess retinal degeneration we used transverse sections stained with DAPI to quantify the thickness of both the ONL and OPL of WT, KO and KO-treated retinas (Fig. 4C). Both the ONL and IPL are thicker in KO-treated retinas compared to KO (Fig. 4Bi), and this was confirmed by quantification of multiple retinas. The KO has both a significantly thinner OPL (Fig. 4Bii) and ONL (Fig. 4Biii) than the KO-treated retinas. The thinner OPL in treated KO mice compared to WT reflects the loss of rod bipolar and horizontal cell profiles, as not all photoreceptors express the target gene with rAAV ^13,19^.

To examine the durability of treatment on retinal function, we compared the ffERGs of the same Control, KO-control, and KO-treated mice at 3 and 14 mpi. Consistent with slow rod degeneration, there is a small but measurable b-wave in KO-control retinas at P35, indicating that there is some transmission from photoreceptor to ON bipolar cells. In the KO-control mice, the b-wave amplitude declines at 3mpi, and no rod-isolated b-wave can be recorded at 14mpi (Fig. 4Di). In contrast, in KO-treated retinas, photoreceptor to bipolar cell transmission is restored, and the restored rod-isolated responses at 3mpi are maintained at 14mpi (Fig. 4Di). We also measured the a-wave of the scotopic ffERG at several flash intensities. At 3 mpi, there were no differences between Control (WT treated rAAV GRK1::GFP), KO-control, and KO-treated, indicating no detectable dysfunction or degeneration (Fig. 4Dii). However, at 14 mpi, the Control a-wave was maintained, but the KO-control declined by ∼50% (Fig. 4Diii). At 14 mpi the a-wave in the KO-treated retinas is increased compared to KO-controls but does not reach the level of Controls (Fig. 4Diii). This is to be expected because not all rods are expected to be infected and express the rAAV.

Together, these results demonstrate that rAAV-mediated α2δ4 expression in KO retinas improves function as measured by the ERG and visual acuity. In addition, it restores the OPL, and reduces rod degeneration, highlighting the neuroprotective potential of this gene therapy approach.

## DISCUSSION

This study investigated whether rAAV-mediated re-expression of the α2δ4 subunit, an auxiliary subunit of the Ca_v_1.4 VDCC, could restore synaptic structure and function of rod photoreceptors of adult *α2δ4^⁻/⁻^* mice. We show that despite the synaptic disruption already present at the time of treatment, restoring α2δ4 expression rescues Ca_v_1.4 channel function, restores the molecular and structural organization of rod ribbon synapses, re-establishes signal transmission from photoreceptors to bipolar cells, and improves visual acuity. The rescue is durable, maintaining retinal structure and function for up to 14 months post-injection. Treatment also slows the rod photoreceptor degeneration otherwise observed in aged *α2δ4^-/-^* mice.

To date, the only clinically approved treatment for an IRD targets an RPE deficiency in Leber’s congenital amaurosis (LCA2), using voretigene neparvovec (Luxturna), which restores retinal function in both adults and children ^11,20^.

In this study, we focus on an iCSNB type caused by mutations in an auxiliary subunit of the Ca_v_1.4 VDCC, which is responsible for glutamate release from photoreceptors ^21^. Ca_v_1.4 is made up of 3 subunits, α1F, β2, and α2δ4, and mutations in any cause iCSNB or rod or cone dystrophies in mouse models ^4,8,22–24^, as do mutations in Ca_v_1.4-modulating proteins such as CaBP4 or ribbon synapse components like RIBEYE and bassoon^23,25^. Because rod photoreceptors make their primary synaptic connection with a single bipolar cell type, the rod bipolar cell, which relays signals via AII amacrine cells to the rest of the retinal circuit, disruption of either glutamate release or its detection is sufficient to cause night blindness. The most common disorder in this group is caused by mutations in the X-linked gene *CACNA1F*, encoding the α1F subunit.

Morphological analyses of iCSNB mouse models reveal a feature that distinguishes them from cCSNB models: complete disassembly of the invaginating photoreceptor synapse, accompanied by ectopic extension of rod bipolar and horizontal cell (HC) processes into the ONL ^15,26,27^. Despite this disorganization, rod axon terminals and their postsynaptic partners may remain in proximity, potentially held together by transsynaptic protein complexes. Some iCSNB models also exhibit slow, progressive retinal degeneration^4^. Critically, gene therapy has not previously been reported for any iCSNB model with this degree of synaptic disorganization, raising the question of whether a synapse this profoundly disrupted can be reformed at all in the mature retina.

Normal rod and cone ribbon synapse formation occurs during development prior to eye opening ^28^. In WT mice, rod bipolar cells and HC axonal processes form an invaginating synapse with rod terminals, the rod spherule, while cone ON bipolar cells and HC dendritic processes form analogous synapses with cones. The invagination process occurs between P8–11 and is initiated when HCs contact the cone axon terminals and is followed by contact with the BCs^28^. In the absence of HC processes, rod bipolar cells fail to invaginate ^16^. In iCSNB models involving knockout of α1F, β2, α2δ4, and CaBP4, the rod spherule fails to develop normally ^4,8,22–24^, as evidenced by loss of the characteristic horseshoe-shaped structure detectable by staining with several antibodies (Figs. 1, 2). The precise mechanism underlying spherule formation remains unresolved but likely reflects both VDCC-mediated calcium influx at the developing synapses and structural transsynaptic interactions.

Although the rod spherule normally forms only during a defined developmental window, we show here that it can be reformed in the adult retina simply by replacing the expression of a single missing component, in our case, α2δ4, in the *α2δ4^-/^*^-^ mouse. This recruits a broad complement of additional synaptic proteins, including LRIT3, mGluR6, TRPM1, GPR179, and PSD95, and is marked by restoration of the horseshoe-shaped structure labeled by RIBEYE, α1F, and α2δ4, mirroring the architecture of WT synapses. Confirming true ultrastructural reformation of the spherule will require further study. Notably, this reformation is not restricted to the OPL: it also occurs at ectopic synapses within the ONL. Most importantly, these reformed synapses are functional; both the ffERG parameters and visual acuity improve following treatment.

Another, important and novel finding of our study is that rAAV-mediated α2δ4 expression in the KO mouse prevents the slow but progressive photoreceptor degeneration that occurs in aged mice. The partial rescue of the ffERG parameters is consistent with the fact that only 40–60% of the retina is infected with the AAV. The mechanism responsible for the degeneration in the KO mice is unknown but is consistent with the emerging concept that synaptic pathology can act as a primary driver of neurodegeneration, and that restoration of synaptic function may have broader neuroprotective effects^29^. The mechanism by which loss of α2δ4 leads to photoreceptor degeneration remains to be fully established, but disruption of calcium channel function and synaptic calcium signaling are likely contributing factors ^9^.

Together, these findings demonstrate, for the first time, that gene therapy can restore synaptic structure and function, improve visual acuity and prevent degeneration in an iCSNB/rod-cone dystrophy model characterized by extensive synaptic disorganization, axonal and dendritic mislocalization, and slow degeneration.

## Acknowledgements

This work was supported by funding from the NIH (R01 EY12354 to R.G.G., M.A.M., N.H., Preston Pope Joyes Endowed Chair in Biochemical Research (R.G.G.) and Kentucky Lions Research in Ophthalmology Endowed Chair (M.A.M.)

## Author Contributions

Conceptualization, R.G.G. and N.H.; Investigation, N.H., M.D; Writing – Original Draft, N.H.; Writing – Review and Editing, R.G.G., M.A.M.; Funding Acquisition, R.G.G., M.A.M. N.H.; Supervision, R.G.G., M.A.M.

## Declaration of Interests

The authors declare no competing interests.

## METHODS

### CONTACT FOR REAGENT AND RESOURCE SHARING

Further information and requests for resources and reagents should be directed to and will be fulfilled by the Lead Contact, Ronald Gregg.

### EXPERIMENTAL MODEL AND SUBJECT DETAILS

All procedures were conducted in accordance with the Society for Neuroscience policies on the use of animals in research and were approved by the University of Louisville Committee for Animal Welfare (UCAW). Animals were housed in the University’s AALAC-approved facility under a 12-hour light/dark cycle. The generation and phenotype of the α2δ4 knockout (KO) mouse line used in this study has been previously described ^15^. Both male and female α2δ4 KO and C57BL/6J (WT) mice were used throughout the study. Mice were anesthetized with intraperitoneal injections of ketamine/xylazine solution (118/11 mg/kg, respectively) diluted in normal Ringer’s solution prior to subretinal injections of rAAVs and ERG recordings. For eye enucleation, mice were euthanized by CO_2_ exposure in accordance with AVMA guidelines. Data from mice with gross retinal damage resulting from the injection procedure were excluded from all analyses.

## METHODS DETAILS

### Antibodies

Antibodies used to label rod and cone synaptic proteins have been previously described and are listed in the key resources table. All antibodies were used at a dilution of 1:1000 and validated for specificity using retinal sections from the respective KO mice.

### rAAV production and subretinal injections

To express wildtype α2δ4 in rod photoreceptors, we used a recombinant adeno-associated virus (rAAV) with gene expression controlled by the rhodopsin (RHO) promoter, which restricts transgene expression to rods. The α2δ4 expression construct was packaged into the rAAV8 capsid by VectorBuilder (Chicago, IL). The rAAV expressing WT α2δ4 under the RHO promoter is designated as rAAV RHO::α2δ4. Subretinal injections were performed by introducing 1 µl of rAAV solution (1 × 10¹³ vg/ml in sterile BSS (balanced salt solution)) into the subretinal space of adult mice (postnatal day 35) using a specialized syringe (www.borghuisinstruments.com). Synaptic protein expression and retinal function were assessed at 5 weeks and 3 and 14 months following rAAV injection.

### Retina preparation for immunohistochemistry

Immunohistochemistry was performed as previously described^12,30^. Mice were euthanized, eyes were enucleated, and the lens was removed. Eyecups were fixed for 20 min in 4% paraformaldehyde in PB (0.1M phosphate buffer, pH 7.4), washed three times in PBS, and cryoprotected in increasing concentrations of sucrose in PB (5%, 10%, and 15% for 1 hr each at room temperature, followed by 20% overnight at 4°C). Eyecups were embedded in a 2:1 OCT/20% sucrose solution and frozen in a liquid nitrogen-cooled isopentane bath. Sections (18 µm) were cut using a Leica 1850 cryostat, mounted on glass slides, and stored at -80°C. Before immunostaining, sections were warmed to 37°C and washed with PBS and PBS containing 0.5% Triton X-100 (PBX) for 5 min each. Sections were blocked in PBX containing 5% normal donkey serum (blocking solution) for 1 hr, then incubated with primary antibody diluted in blocking solution overnight at room temperature. Sections were washed four times for 10 min each in PBX, incubated with secondary antibody (1:1000) in PBX for 1 hr at room temperature, washed three times in PBX and once in PBS for 10 min each, and coverslipped using Vectashield (Vector Laboratories, Burlingame, CA). Images were acquired using an Olympus FV4000 confocal microscope and uniformly adjusted for brightness using CellSens software (Olympus). ONL and OPL thickness was measured in retinal sections stained with DAPI using FIJI/ImageJ software. The scale was set based on the pixel size from the confocal microscope settings. For each image, five thickness measurements were taken at evenly spaced locations across the retina. Three images were analyzed per retina, and 3-4 mice per group were included in the analysis.

### Electroretinography

ERG recordings were performed as previously described^31^. Mice were dark-adapted overnight and anesthetized with a ketamine/xylazine solution (118/11 mg/kg) diluted in Ringer’s solution. All preparations were performed under dim red light to minimize light exposure. Pupils were dilated with topical applications of 0.625% phenylephrine hydrochloride and 0.25% tropicamide, and the corneal surface was anesthetized with 1% proparacaine HCl. Body temperature was maintained throughout the procedure using a feedback-controlled electric heating pad (TC1000; CWE Inc.). A contact lens with a gold electrode (LKC Technologies Inc.) was placed on the cornea and moistened with artificial tears (Tears Again; OCuSOFT, Gaithersburg, MD) to ensure proper electrical contact. Ground and reference needle electrodes were positioned in the tail and on the midline of the forehead, respectively.

Scotopic responses were recorded from dark-adapted mice using test flashes ranging from -3.6 to 1.4 log cd·s/m². Photopic responses were recorded after 5 min of light adaptation to a rod-saturating background of 20 cd/m², using test flashes ranging from -0.8 to 1.4 log cd·s/m².

### Evaluation of Visual Acuity

Visual acuity was measured using a two-alternative forced choice method ^18^, modified to measure both photopic and scotopic visual acuity^17^. The task has three phases: molding, training, and testing. In the molding phase, mice are acclimatized to the water and learn to escape to a hidden platform. In the training phase, mice learn that the grating on the monitor is associated with the hidden platform and complete the task successfully in 8 out of 10 trials. In the testing phase, the spatial frequency of the grating is systematically increased to determine the threshold at which the mouse can no longer discriminate the grating from the equiluminant side. All groups, WT, *α2δ4^-/-^*, and rAAV RHO::α2δ4-treated *α2δ4^-/-^* mice were first trained and tested under photopic conditions with screens set to 100 cd/m². Mice were then dark-adapted for 60 min at 0.01 cd/m² before scotopic visual acuity was measured with both screens set to 0.025 cd/m².

### Statistical Analysis

Statistical analysis was performed using Prism 11.0.0 (GraphPad Software, La Jolla, CA). Details of the statistical tests used are provided in the text and figure legends. Where multiple comparisons were made, Tukey post-hoc tests were applied and adjusted p-values (p_adj_) are reported. Statistical significance was defined as p_adj_≤ 0.05.

## KEY RESOURCES TABLE

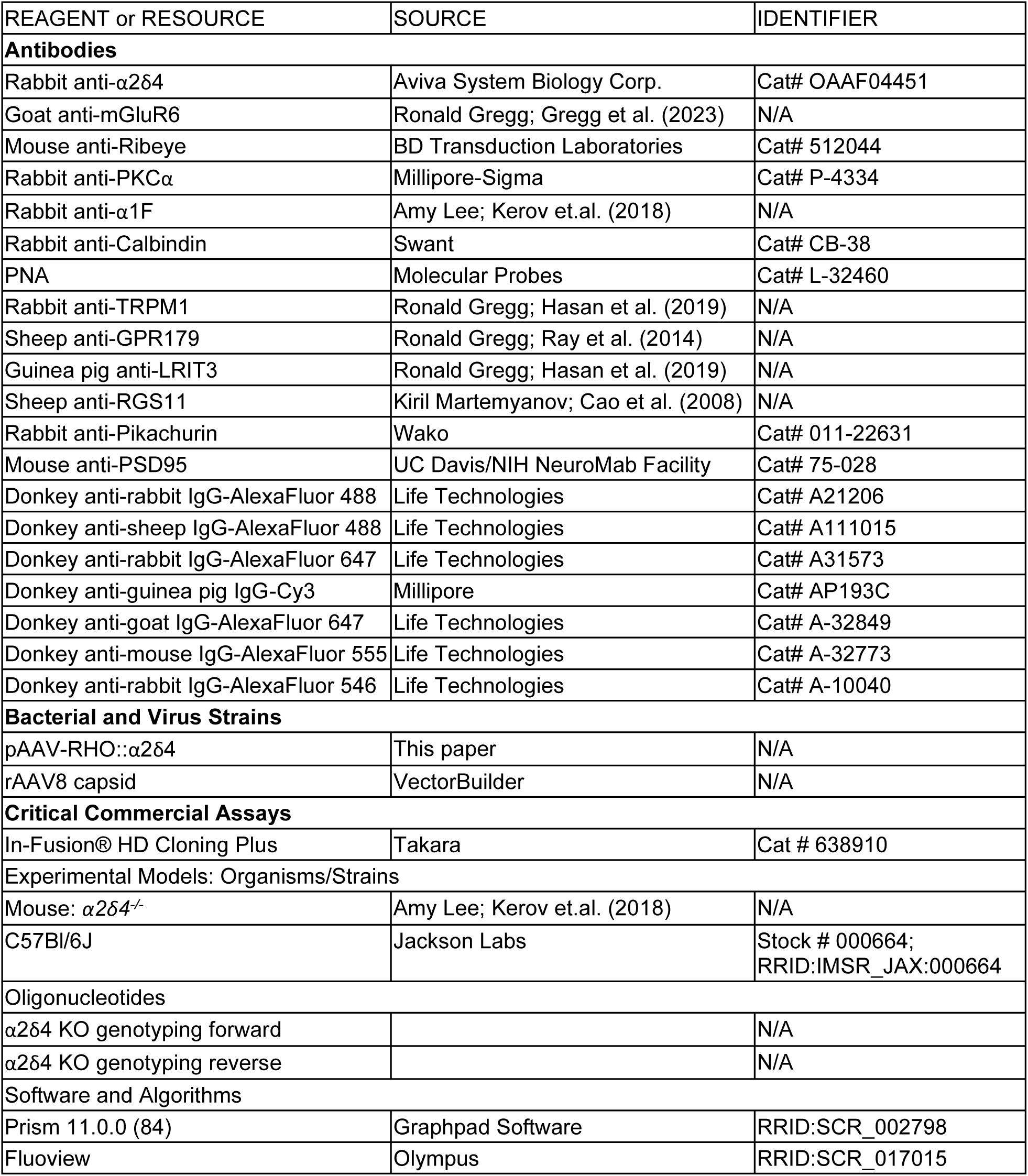

| REAGENT or RESOURCE | SOURCE | IDENTIFIER |
| --- | --- | --- |
| **Antibodies** |  |  |
| Rabbit anti-$\alpha 2\delta 4$ | Aviva System Biology Corp. | Cat# OAAF04451 |
| Goat anti-mGluR6 | Ronald Gregg; Gregg et al. (2023) | N/A |
| Mouse anti-Ribeye | BD Transduction Laboratories | Cat# 512044 |
| Rabbit anti-PKCα | Millipore-Sigma | Cat# P-4334 |
| Rabbit anti-$\alpha 1F$ | Amy Lee; Kerov et.al. (2018) | N/A |
| Rabbit anti-Calbindin | Swant | Cat# CB-38 |
| PNA | Molecular Probes | Cat# L-32460 |
| Rabbit anti-TRPM1 | Ronald Gregg; Hasan et al. (2019) | N/A |
| Sheep anti-GPR179 | Ronald Gregg; Ray et al. (2014) | N/A |
| Guinea pig anti-LRIT3 | Ronald Gregg; Hasan et al. (2019) | N/A |
| Sheep anti-RGS11 | Kiril Martemyanov; Cao et al. (2008) | N/A |
| Rabbit anti-Pikachurin | Wako | Cat# 011-22631 |
| Mouse anti-PSD95 | UC Davis/NIH NeuroMab Facility | Cat# 75-028 |
| Donkey anti-rabbit IgG-AlexaFluor 488 | Life Technologies | Cat# A21206 |
| Donkey anti-sheep IgG-AlexaFluor 488 | Life Technologies | Cat# A111015 |
| Donkey anti-rabbit IgG-AlexaFluor 647 | Life Technologies | Cat# A31573 |
| Donkey anti-guinea pig IgG-Cy3 | Millipore | Cat# AP193C |
| Donkey anti-goat IgG-AlexaFluor 647 | Life Technologies | Cat# A-32849 |
| Donkey anti-mouse IgG-AlexaFluor 555 | Life Technologies | Cat# A-32773 |
| Donkey anti-rabbit IgG-AlexaFluor 546 | Life Technologies | Cat# A-10040 |
| **Bacterial and Virus Strains** |  |  |
| pAAV-RHO::$\alpha 2\delta 4$ | This paper | N/A |
| rAAV8 capsid | VectorBuilder | N/A |
| **Critical Commercial Assays** |  |  |
| In-Fusion® HD Cloning Plus | Takara | Cat # 638910 |
| **Experimental Models: Organisms/Strains** |  |  |
| Mouse: $\alpha 2\delta 4^{-/-}$ | Amy Lee; Kerov et.al. (2018) | N/A |
| C57Bl/6J | Jackson Labs | Stock # 000664;<br>RRID:IMSR_JAX:000664 |
| **Oligonucleotides** |  |  |
| $\alpha 2\delta 4$ KO genotyping forward | | N/A |
| $\alpha 2\delta 4$ KO genotyping reverse | | N/A |
| **Software and Algorithms** |  |  |
| Prism 11.0.0 (84) | Graphpad Software | RRID:SCR_002798 |
| Fluoview | Olympus | RRID:SCR_017015 |

## Notes

### Competing Interest Statement

The authors have declared no competing interest.

