## Supplemental Figures for "Therapeutic restoration of synaptic architecture, retinal and visual function, and prevention of retinal degeneration in a mouse model of retinal dystrophy"

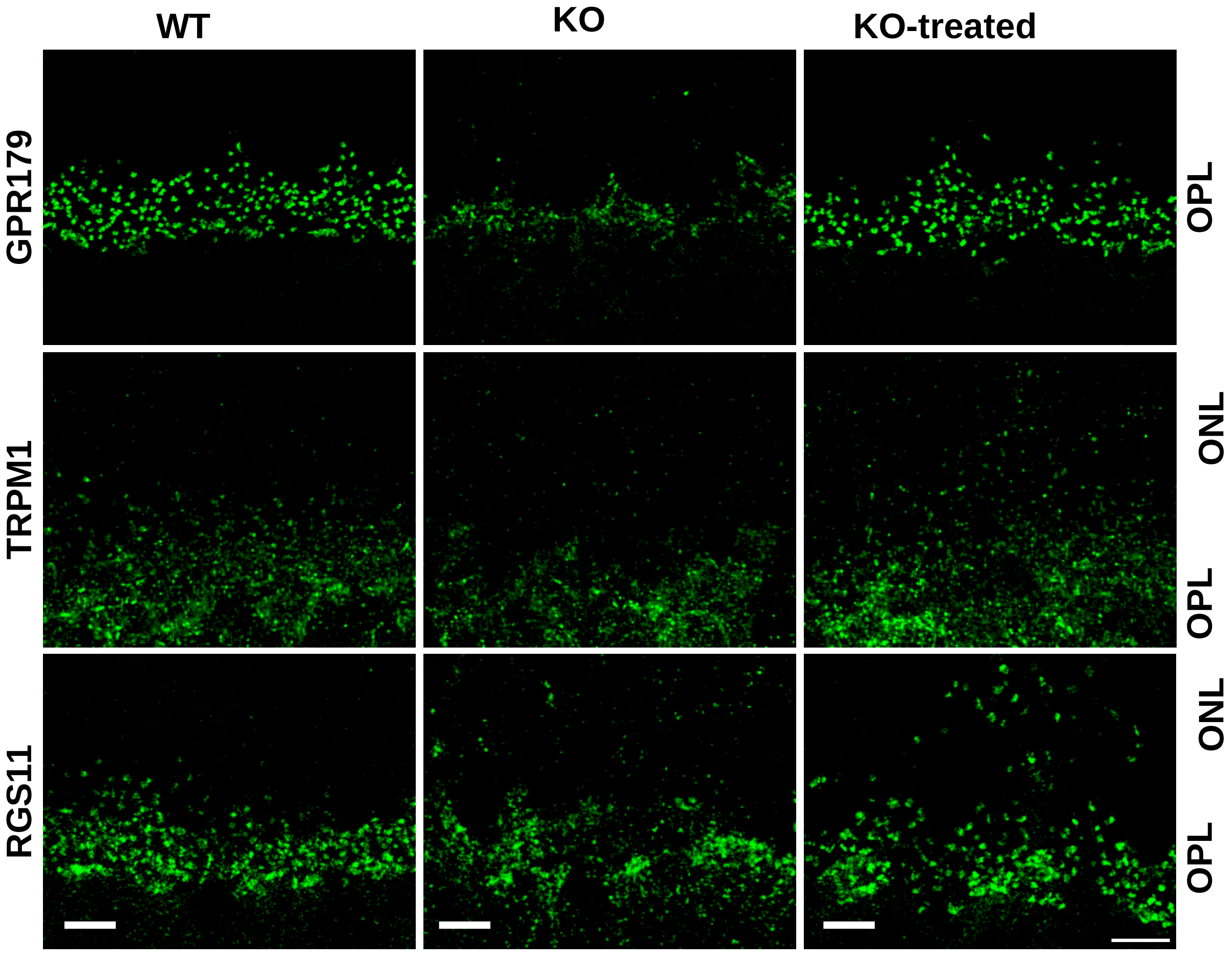


**Figure S1. Expression of α2δ4 in rods restores post-synaptic components.** Representative images of transverse retinal sections of WT, KO and KO-treated retinas stained with antibodies to GPR179, TRPM1, and RGS11. Images are representative of those from at least 5 retinas. Scale bars = 5µm. Staining is restricted to the OPL in WT, and decreased and localized throughoutot the OPL and ONL in the KO. In the KO-treated retina the puncta in the OPL are more robust as are those in the ONL.

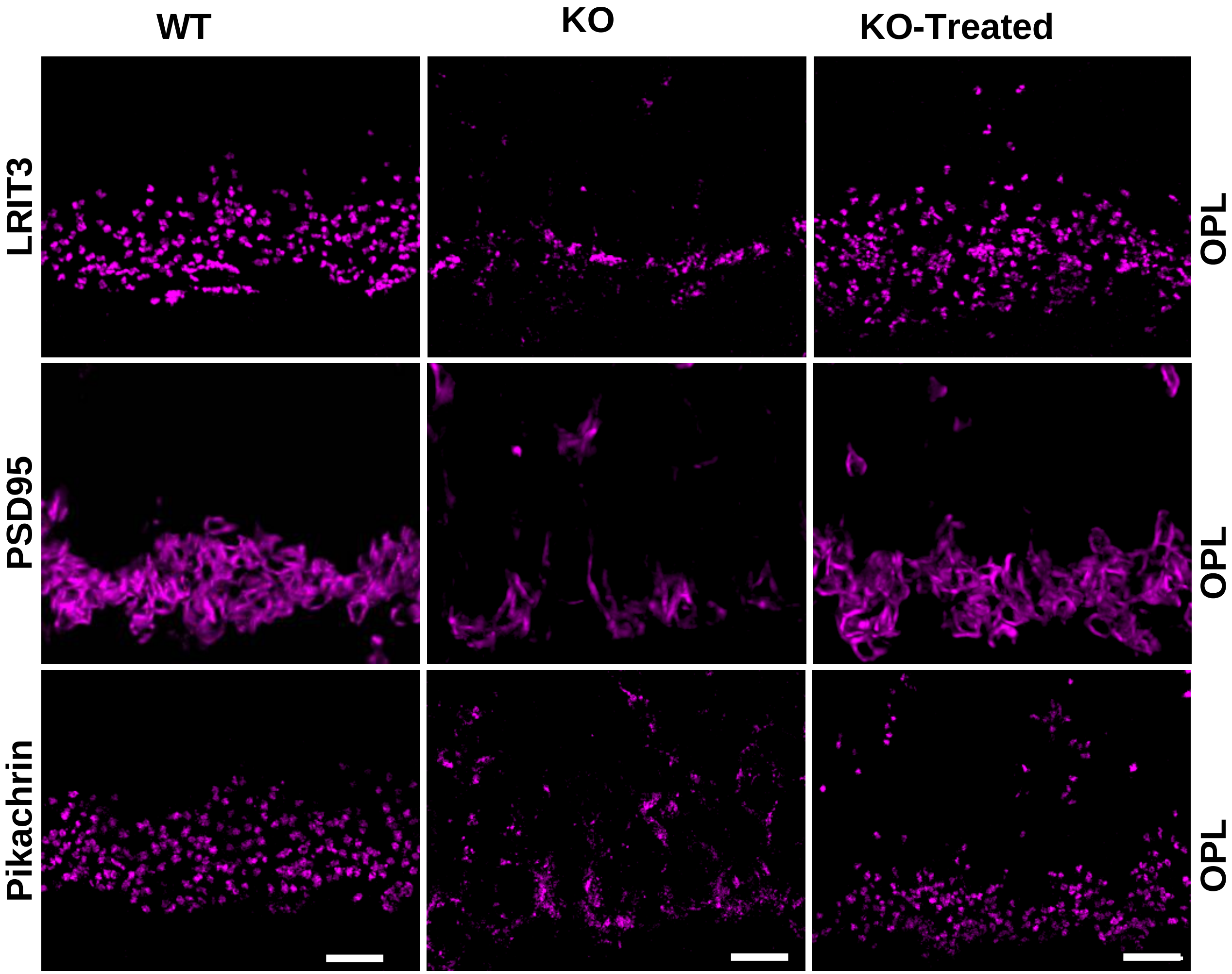


**Figure S2. Expression of α2δ4 in rods restores pre-synaptic components.** Representative images of transverse retinal sections of WT, KO and KO-treated retinas stained with antibodies to LRIT3, PSD95 and pikachurin. Images are representative of those from at least 5 retinas. Scale bars = 5μm. Staining is restricted to the OPL in WT, and deacreased and localized throughout the OPL and ONL in KO. In the KO-treated retinas the puncta in the OPL are more robust as are those in the ONL.

### Figure S3

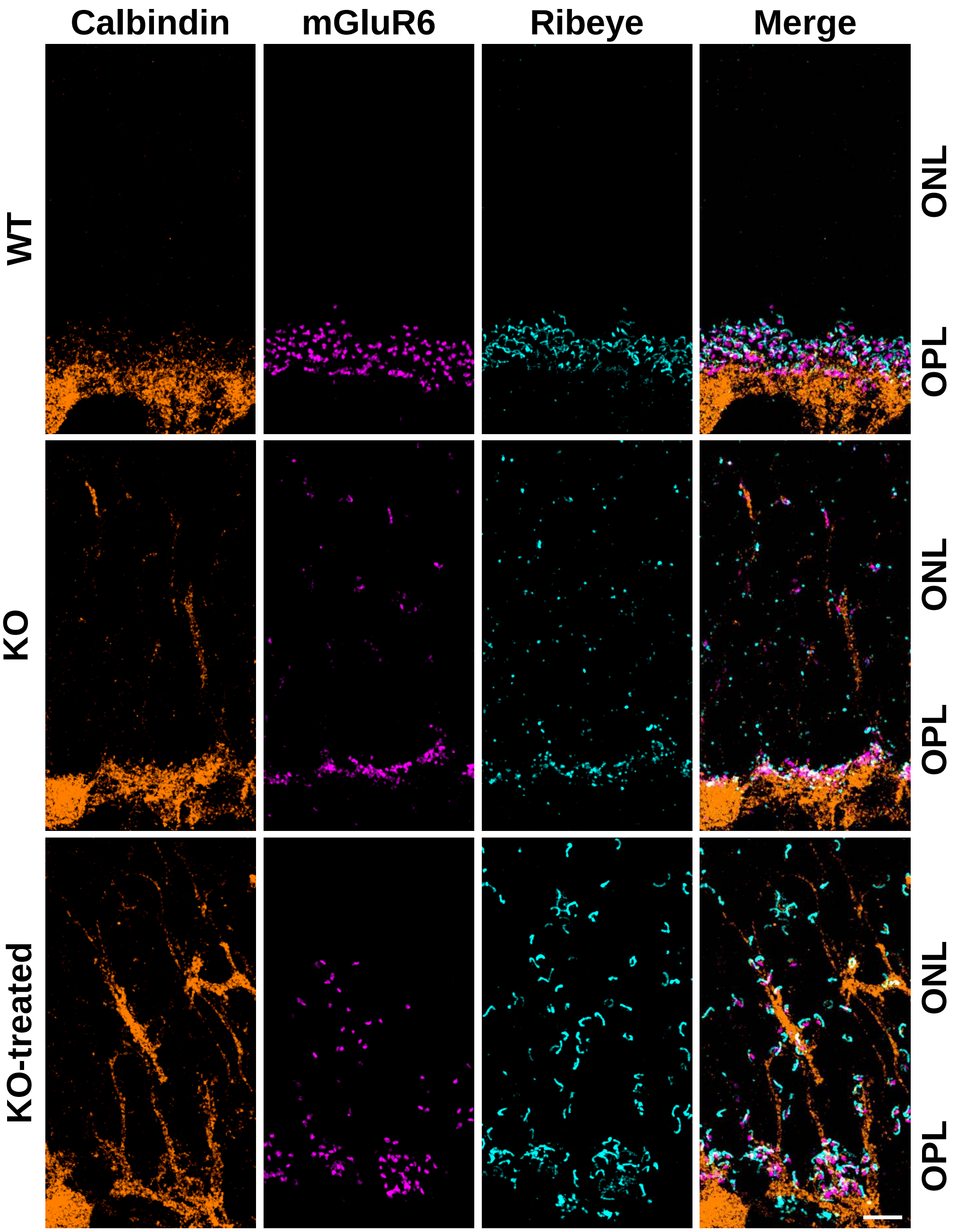


**Figure S3. Expression of α2δ4 in rods restores restores rod synaptic organization.** Representative images of transverse retinal sections of WT, KO and KO-treated retinas stained with antibodies to calbindin, mGluR6 and ribeye. Images are representative of those from at least 5 retinas. Scale bars = 5μm. Staining  is restricted to the OPL in WT, and ribeye punctate staining localized throughout the OPL and ONL in KO. In the KO-treated retinas ribeye horseshoe-shaped structures are reformed in the OPL as well as in the ONL.
